# Nitrous oxide reduction by two partial denitrifying bacteria requires denitrification intermediates that cannot be respired

**DOI:** 10.1101/2022.06.21.497020

**Authors:** Breah LaSarre, Ryan Morlen, Gina C. Neumann, Caroline S. Harwood, James B. McKinlay

## Abstract

Denitrification is a form of anaerobic respiration wherein nitrate (NO_3_^-^) is sequentially reduced via nitrite (NO_2_^-^), nitric oxide, and nitrous oxide (N_2_O) to dinitrogen gas (N_2_) by four reductase enzymes. Partial denitrifying bacteria possess only one, or some, of these four reductases and use them as independent respiratory modules. However, it is unclear if partial denitrifiers sense and respond to denitrification intermediates outside of their reductase repertoire. Here we tested the denitrifying capabilities of two purple nonsulfur bacteria, *Rhodopseudomonas palustris* CGA0092 and *Rhodobacter capsulatus* SB1003. Each had denitrifying capabilities that matched their genome annotation; CGA0092 reduced NO_2_^-^ to N_2_ and SB1003 reduced N_2_O to N_2_. For each bacterium, N_2_O reduction could be used for both electron balance during growth on electron-rich organic compounds in light and for energy transformation via respiration in the dark. However, N_2_O reduction required supplementation with a denitrification intermediate, including those for which there was no associated denitrification enzyme. For CGA0092, NO_3_^-^ served as a stable, non-catalyzable molecule that was sufficient to activate N_2_O reduction. Using a β-galactosidase reporter we found that NO_3_^-^ acted, at least in part, by stimulating N_2_O reductase gene expression. In SB1003, NO_2_^-^, but not NO_3_^-^, activated N_2_O reduction but NO_2_^-^ was slowly removed, likely by a promiscuous enzyme activity. Our findings reveal that partial denitrifiers can still be subject to regulation by denitrification intermediates that they cannot use.

**Importance:** Denitrification is a form of microbial respiration wherein nitrate is converted via several nitrogen oxide intermediates into harmless dinitrogen gas. Partial denitrifying bacteria, which individually have some but not all denitrifying enzymes, can achieve complete denitrification as a community by cross-feeding nitrogen oxide intermediates. However, the last intermediate, nitrous oxide (N_2_O), is a potent greenhouse gas that often escapes, motivating efforts to understand and improve the efficiency of denitrification. Here we found that at least some partial denitrifying N_2_O reducers can sense and respond to nitrogen oxide intermediates that they cannot otherwise use. The regulatory effects of nitrogen oxides on partial denitrifiers are thus an important consideration in understanding and applying denitrifying bacterial communities to combat greenhouse gas emissions.

## Introduction

Denitrification is a multistep respiratory pathway that sequentially reduces nitrate (NO_3_^-^) via nitrite (NO_2_^-^), nitric oxide (NO), and nitrous oxide (N_2_O) to dinitrogen gas (N_2_) (1, 2) (Fig. 1A). Denitrifying bacteria are important in several contexts. Denitrifiers in the human gut help fight pathogens and maintain vascular homeostasis through the generation of NO_2_^-^ and NO (3). Denitrification is also important to the global nitrogen cycle, returning nitrogen to the atmosphere as N_2_. However, N_2_O often escapes denitrifying communities before it can be reduced to N_2_. N_2_O is a potent greenhouse gas that damages the ozone layer (4). N_2_O emissions have increased to concerning levels, primarily due to transformation of NO_3_^-^ in agricultural fertilizers to N_2_O by naturally occurring denitrifying bacteria in the soil (5, 6). Thus, there is a need to better understand and improve the efficiency of denitrification.

**Fig. 1.**
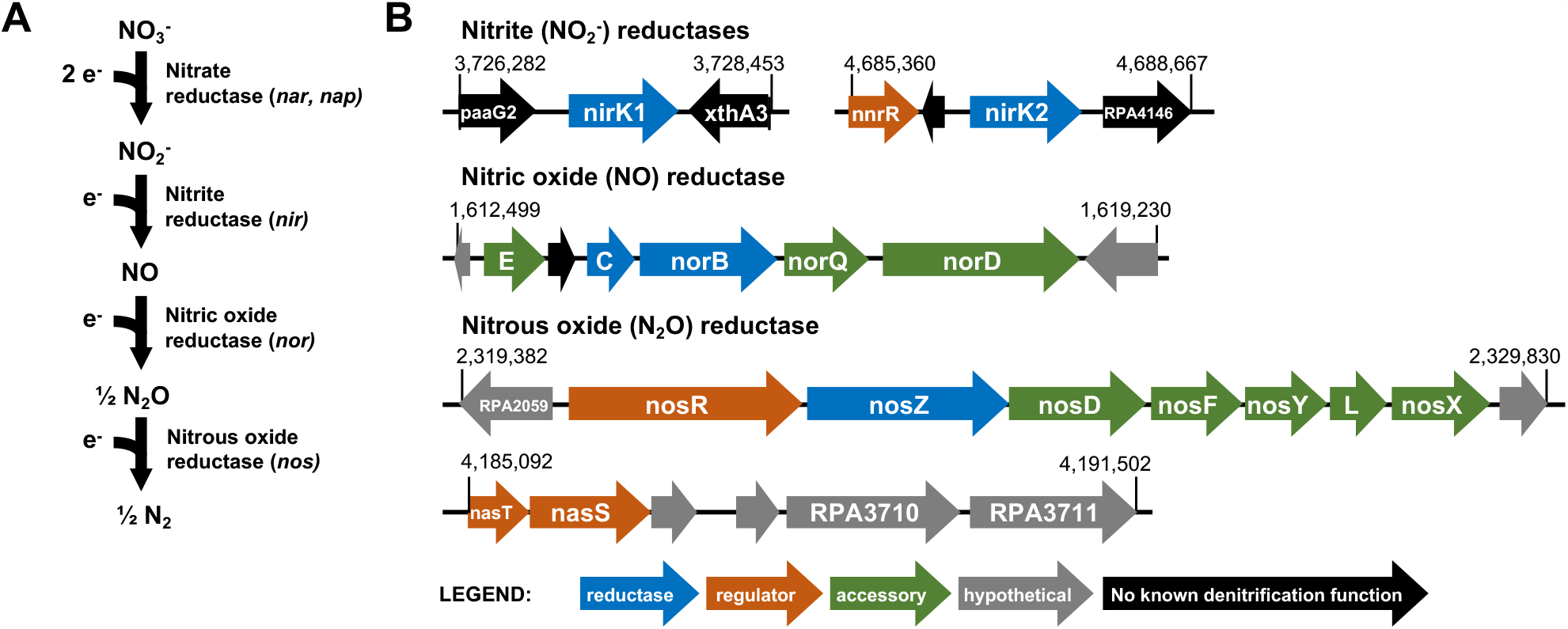
General denitrification pathway (A) and denitrification genes annotated in *R. palustris* CGA0092. **(B)**. Numbers indicate the chromosome nucleotide positions. Several CRP/Fnr-family transcriptional regulators with >25% sequence identity to known denitrification regulators are not shown.

Many bacteria lack a complete denitrification pathway and are thus called partial or truncated denitrifiers (7-11). Partial denitrifiers use single or multiple steps of the pathway as independent respiratory modules (1, 2). Although incapable of reducing NO_3_^-^ to N_2_ on their own, partial denitrifiers are important contributors to complete denitrification as a community process, with intermediates cross-fed between community members that have different segments of the pathway (7-12). Notably, nitrogen oxides (NO_3_^-^, NO_2_^-^, NO, or N_2_O) not only serve as substrates for denitrification reductases can also act as regulators of denitrification. Although regulatory roles have been well-characterized in bacteria capable of complete denitrification, regulatory roles in partial denitrifiers have received comparatively less attention. In particular, it is unclear if the regulatory effects of nitrogen oxides in partial denitrifiers matches their reductase repertoire.

Here we characterized the ability of two purple non-sulfur bacteria (PNSB) that are putative partial denitrifiers, *Rhodopseudomonas palustris* CGA0092 and *Rhodobacter capsulatus* SB1003, to carry out denitrification under photoheterotrophic and chemoheterotrophic conditions. Under phototrophic conditions, where light is the energy source, we tested if nitrogen oxides can serve as an essential electron acceptor to maintain electron balance during growth on the electron-rich substrate butyrate. Under chemotrophic conditions, we tested if nitrogen oxides can serve as an essential electron acceptor to generate energy via oxidative phosphorylation. As expected, each bacterium was only able to grow using the nitrogen oxides for which corresponding reductases were annotated in their genomes. However, N_2_O utilization required supplementation with additional nitrogen oxides other than N_2_O, including nitrogen oxides for which there was no predicted reductase and which did not support growth on their own. Our results indicate that at least some partial denitrifiers require nitrogen oxides that they cannot respire to reduce N_2_O.

## Results

### *R. palustris* CGA0092 has a partial denitrification pathway

*R. palustris* is one of the most metabolically versatile PNSB (13), yet little is known about its ability to respire anaerobically. Unlike some other model PNSB, CGA009, and its derivative CGA0092 (14) used herein, cannot grow via respiration with dimethylsulfoxide (15-17). However, according to its genome sequence, it should be capable of partial denitrification, as it has putative enzymes for converting NO_2_^-^ to N_2_ (10, 13) (Fig. 1B). Expanding on past analyses by others (10, 13), we used PSI-BLAST to verify that there are no genes with significant similarity to *nar, nap*, or *nas* nitrate reductase genes nor to *nasB, nirB, nirD, nirA, nrf*, or eight-heme nitrite reductase genes (Table S1) (18).

When incubated anaerobically in darkness, PNSB typically use electron acceptors to establish a proton motive force and generate ATP. When incubated in light, PNSB generate ATP by photophosphorylation but electron acceptors, such as CO_2_ or NaHCO_3_, are required for growth on electron-rich compounds like butyrate to prevent an accumulation of reduced electron carriers that halts metabolism; butyrate contains more electrons than can be incorporated into biomass and so the excess electrons must be deposited on an electron acceptor or released as H_2_ (19, 20). Given that *R. palustris* grows best in light, we first examined if it could use denitrification intermediates as electron acceptors during growth with butyrate.

In agreement with the apparent lack of NO_3_^-^ reductase in the CGA0092 genome (Fig. 1B), phototrophic growth on butyrate was not supported when supplemented with a wide range of NaNO_3_ concentrations (Fig. 2A). However, growth was observed when CGA0092 was provided with 1mM NaNO_2_ (Fig. 2A). We determined that this concentration was near the toxicity limit because it caused a lag in phototrophic growth on succinate, which does not require supplementation with an electron acceptor (Fig. 2B). We verified NO_2_^-^ utilization using the colorimetric Griess assay. All of the NO_2_^-^ removed could be accounted for in the accumulated N_2_ and N_2_O, as measured by gas chromatography (Fig. 2C). N_2_ and N_2_O levels were, in fact, higher than what could be explained by conversion of the supplied NO_2_^-^, although not significantly different from the expected 1:1 correspondence. If real, the excess nitrogen was likely due to contamination with atmospheric N_2_ during sampling and an inability to distinguish N_2_O from CO_2_ produced by other metabolic reactions (CO_2_ and N_2_O coeluted in our gas chromatography method).

**Fig. 2.**
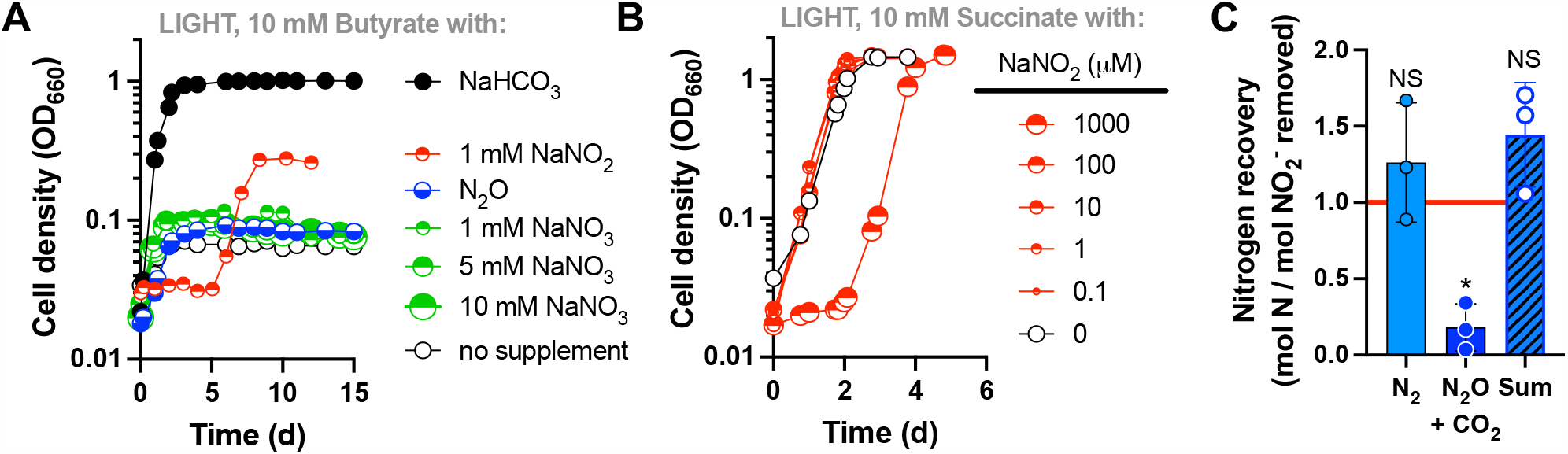
NaNO_2_ supports phototrophic growth of CGA0092 on butyrate within toxicity limits. **A**. Phototrophic growth with butyrate and various potential electron acceptors. Single representatives are shown. Similar trends were observed for three biological replicates except for conditions exploring different NaNO_3_ concentrations where only single representatives were used. **B**. Phototrophic growth with succinate, a condition that readily supports growth without supplementation with an electron acceptor, with various concentrations of NaNO_2_ to identify the toxicity limit. Cultures had a 100% Ar headspace unless N_2_O is indicated (100% N_2_O). **C**. Proportion of NO_2_^-^ recovered as N_2_ and N_2_O. All of the supplied NO_2_^-^ was removed. N_2_O peak area includes a minor contribution of CO_2_ due to coelution during gas chromatography. Each point represents an independent biological replicate. *, significantly different from 1 (p < 0.05); NS, not significantly different from 1.0 (p > 0.05), determined by a one sample t and Wilcoxon test.

Generation of N_2_ from NO_2_^-^ indicated that the latter three reductases for denitrification are active in CGA0092. We did not directly address reduction of exogenously added NO because it is highly toxic and would likely be impossible to add in amounts that would be practical to yield observable growth. We directly addressed N_2_O reduction by providing CGA0092 with a headspace of 100% N_2_O. However, no growth was observed within 15 days (Fig. 2A), suggesting that N_2_O alone cannot activate N_2_O reduction.

### Photoheterotrophic N_2_O reduction by CGA0092 requires NaNO_2_ or NaNO_3_

In some bacteria, denitrification intermediates other than N_2_O enhance, or are required for, N_2_O reduction (1, 2, 21-23). For example, NO_2_^-^ induces N_2_O reductase at the transcriptional level via the regulatory protein NnrR, though NO_2_^-^ might first need to be converted to NO (23). NO_3_^-^ induces N_2_O reductase via NasTS regulatory proteins and an anti-terminator mechanism that affects transcription of *nos* genes (24, 25). CGA0092 encodes NnrR upstream of NirK2 and NasTS homologs upstream of a gene cluster encoding a potential nitrite/sulfite reductase (RPA3710-11; Fig. 1B). We thus tested if NaNO_2_ or NaNO_3_ could enable growth with N_2_O. In agreement with our hypothesis, micromolar amounts of NaNO_2_ as low as 1 µM stimulated growth with N_2_O, with final cell densities increasing in accordance with the amount of NaNO_2_ added (Fig. 3A). NaNO_2_ at 100 µM caused a 4-day lag in growth (Fig. 3A), suggesting that this level of NO_2_^-^ was slightly toxic under these conditions. Notably, the increase in final cell density afforded by high amounts of NaNO_2_ depended on the presence of N_2_O, as growth with 100 µM NaNO_2_ alone was much lower (Fig. 3A). This indicated that N_2_O was being used as the primary electron acceptor in the presence of NO_2_^-^, despite that N_2_O could not serve as an electron acceptor when provided alone (Fig. 2A). We speculate that exhaustion of NO_2_^-^ eliminated the activation of N_2_O reduction, thereby eliminating the ability to use N_2_O as an electron acceptor; consequently, growth on butyrate with N_2_O lasts only as long as the pool of NaNO_2_.

**Fig. 3.**
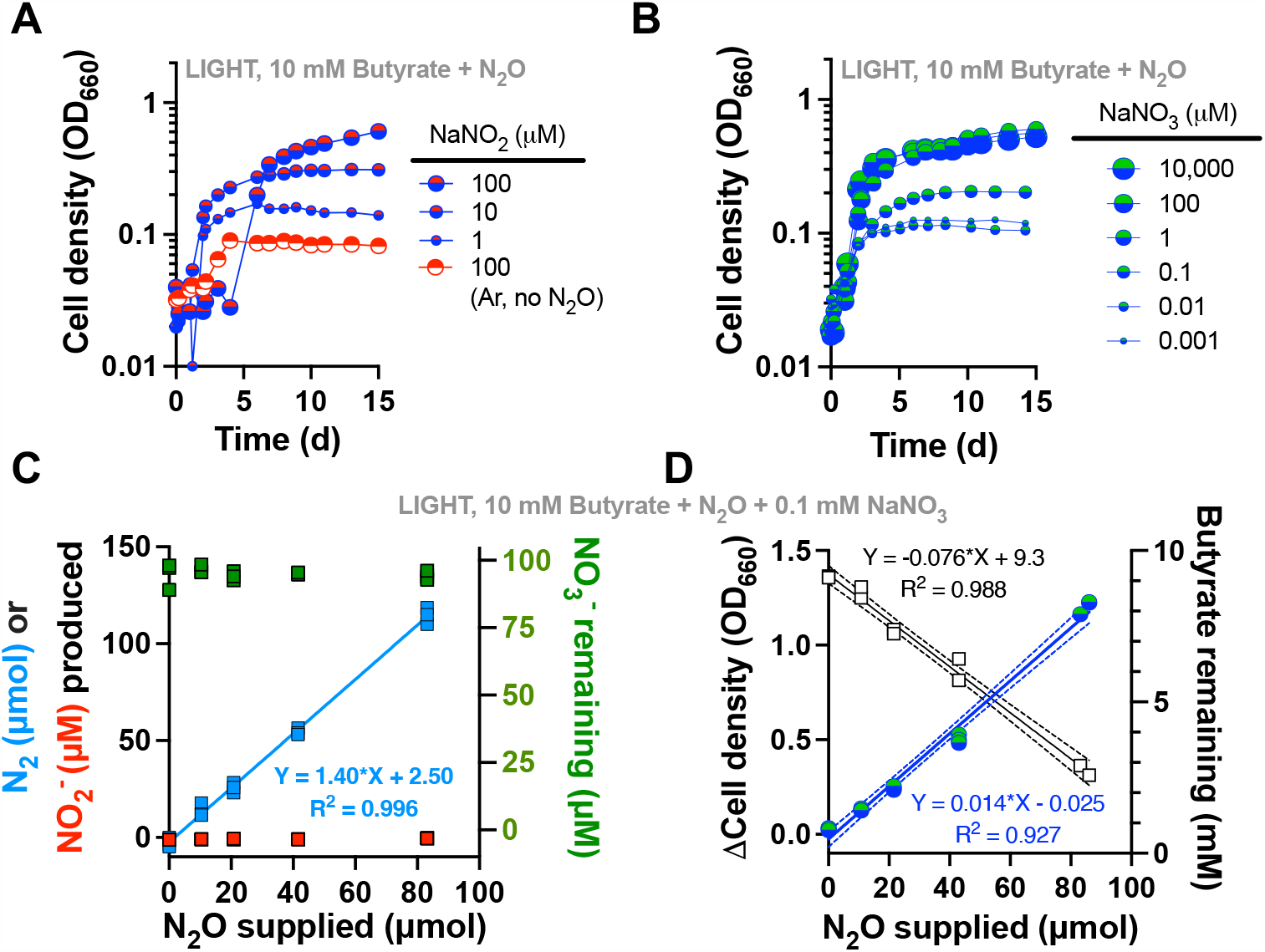
NaNO_2_ or NaNO_3_ is required for phototrophic N_2_O utilization by CGA0092. **A**. Phototrophic growth with butyrate +/- N_2_O and different concentrations of NaNO_2_. The ‘Ar, no N_2_O’ control had a 100% argon headspace. **B**. Phototrophic growth with butyrate + N_2_O and different concentrations of NaNO_3_. Single representatives were surveyed. **C, D**. Changes in N_2_, NO_3_^-^, NO_2_^-^, cell density, and butyrate in N_2_O-limited cultures. Measurements were taken at inoculation and when the maximum cell density was reached. Each point represents a single independent culture. All N_2_O was removed by stationary phase. Linear regression for NO_3_^-^ (**C**) gave a slope that was not significantly different from zero (p = 0.57).

Despite lacking NO_3_^-^ reductase, NaNO_3_ was also sufficient to stimulate phototrophic growth on butyrate with N_2_O (Fig. 3B). Similar growth trends were observed between 1 µM to 10 mM NaNO_3_, indicating that NaNO_3_ is relatively non-toxic and suggesting that NO_3_^-^ was not being reduced. Indeed, NO_3_^-^ levels were stable when we used 0.1 mM NaNO_3_ to stimulate photoheterotrophic N_2_O reduction (Fig. 3C). The amount of N_2_O provided, all of which was ultimately removed, was linearly correlated with N_2_ generated (Fig. 3C), although we again observed more N_2_ generated than should be possible from the N_2_O provided, likely due to contamination with atmospheric N_2_. Culture growth and butyrate consumed were also linearly correlated with N_2_O supplied (and removed), further demonstrating the use of N_2_O as an electron acceptor (e.g., in place of NaHCO_3_; Fig. 2A) for phototrophic growth with butyrate.

To determine if the requirement of NaNO_3_ or NaNO_2_ for N_2_O reduction is manifested at the level of N_2_O reductase expression (the combination of transcription and translation), we created a reporter that fused the region upstream of *nosR*, which should contain both a transcriptional promoter and a ribosomal binding site, to the *lacZ* gene at the start codon and integrated the reporter into the CGA0092 chromosome. We then grew the resulting reporter strain (CGA4070) under phototrophic conditions with acetate, a condition where an electron acceptor supplement is not required, and added either NaCl (negative control), NaNO_3_, NaNO_2_, or N_2_O as possible inducers of expression. In agreement with growth trends (Fig. 3), NaNO_3_ and NaNO_2_, but not N_2_O, led to a significant, albeit low (1.6 – 2.4-fold), increase in LacZ activity over the NaCl control (Fig. 4). Thus, NaNO_3_ and NaNO_2_ are inducers of *nos* gene expression, although the relatively small effect suggests that there might be additional levels of regulatory control.

**Fig. 4.**
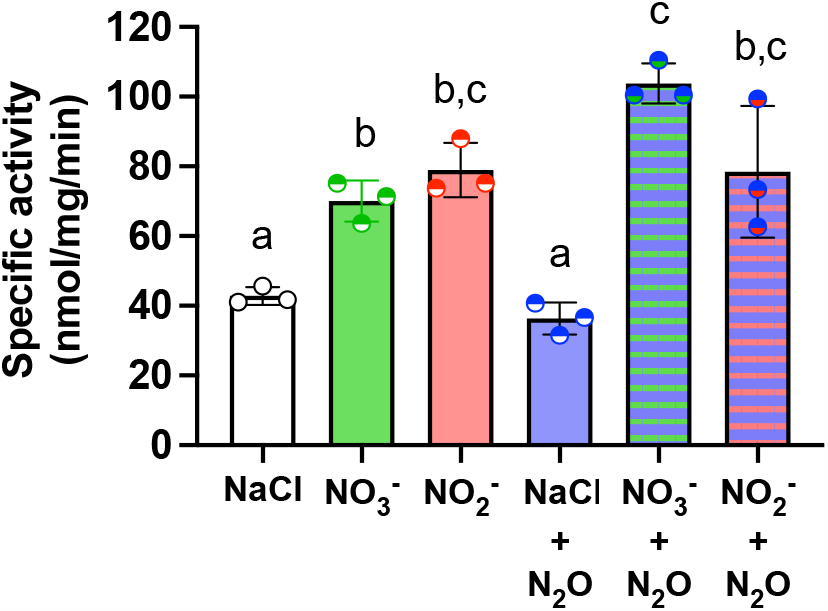
NaNO_3_ and NaNO_2_, but not N_2_O, positively affect *nosR* expression. β-galactosidase measurements were made in cell extracts of CGA4070, which harbors a chromosomally integrated *nosR* promoter*–lacZ* fusion. CGA4070 was grown phototrophically with 23 mM acetate and 0.1 mM NaCl, NaNO_3_ or NaNO_2_. Cultures received 4 ml N_2_O, where indicated. Each point represents a single independent culture. All values were corrected for a background *o*-nitrophenol production rate of 8.7 nmol/mg/min as measured in cell extracts from CGA0092 grown under identical conditions with 0.1 mM NaCl or NaNO_3_; activity was not significantly different between conditions with NaCl or NaNO_3_. Floating letters indicate significant differences between strains (One-way ANOVA with Tukey post-test)p < 0.5).

### N_2_O plus NaNO_3_ can rescue photoheterotrophic growth of *R. palustris* Calvin cycle mutants

Under most photoheterotrophic growth conditions, the CO_2_-fixing Calvin cycle is essential to maintain electron balance, even on relatively oxidized substrates like succinate (19, 26, 27). To distinguish this essential electron balancing role from the Calvin cycle’s better known role in carbon assimilation, alternative electron acceptors are a useful tool because they permit growth of Calvin cycle mutants (28, 29). Thus far, the only method known to allow growth of *R. palustris* Calvin cycle mutants under conditions where the cycle is normally essential was via NifA* mutations that result in constitutive nitrogenase activity (19, 26, 27). NifA* mutants dispose of excess electrons as H_2_, an obligate product of the nitrogenase reaction. However, our results suggested that N_2_O could be used as an electron acceptor to grow *R. palustris* Calvin cycle mutants without additional genetic intervention. Indeed, N_2_O with NaNO_3_ rescued an *R. palustris* Calvin cycle mutant (ΔCalvin) during phototrophic growth on succinate (Fig. 5). N_2_O reduction resulted in more immediate growth than a NifA* mutation. However, growth eventually slowed and the culture reached a lower final cell density that the ΔCalvin NifA* mutant (Fig. 5). Gas chromatographic analysis of headspace samples confirmed that growth of cultures with N_2_O plus NaNO_3_ was not due to spontaneous mutations that enabled H_2_ production (i.e., no H_2_ was detected; data not shown).

**Fig. 5.**
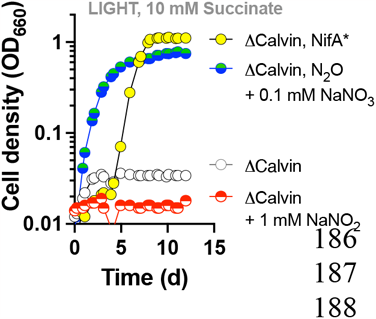
N_2_O plus NaNO_3_ supports phototrophic growth of an *R. palustris* Calvin cycle deletion mutant. Cultures had a 100% Ar headspace unless N_2_O is indicated (100% N_2_O). Single representatives are shown. Similar trends were observed for three biological replicates. ΔCalvin is strain CGA4008; ΔCalvin, NifA* is strain CGA4011.

### NaNO_3_ does not improve *R. palustris* photoheterotrophic growth with NaNO_2_

We wondered if NaNO_3_ might also improve growth with NO_2_^-^, perhaps by stimulating NO_2_^-^ reductase activity. However, supplementation with NaNO_3_ did not affect photoheterotrophic growth trends on butyrate with 1 mM NaNO_2_, even when NaNO_3_ was also added to starter cultures as a possible ‘pre-inducing’ condition (Fig. 6A). The same strategy also did not decrease the lag phase during phototrophic growth on succinate with 1 mM NaNO_2_ (Fig. 6B).

**Fig. 6.**
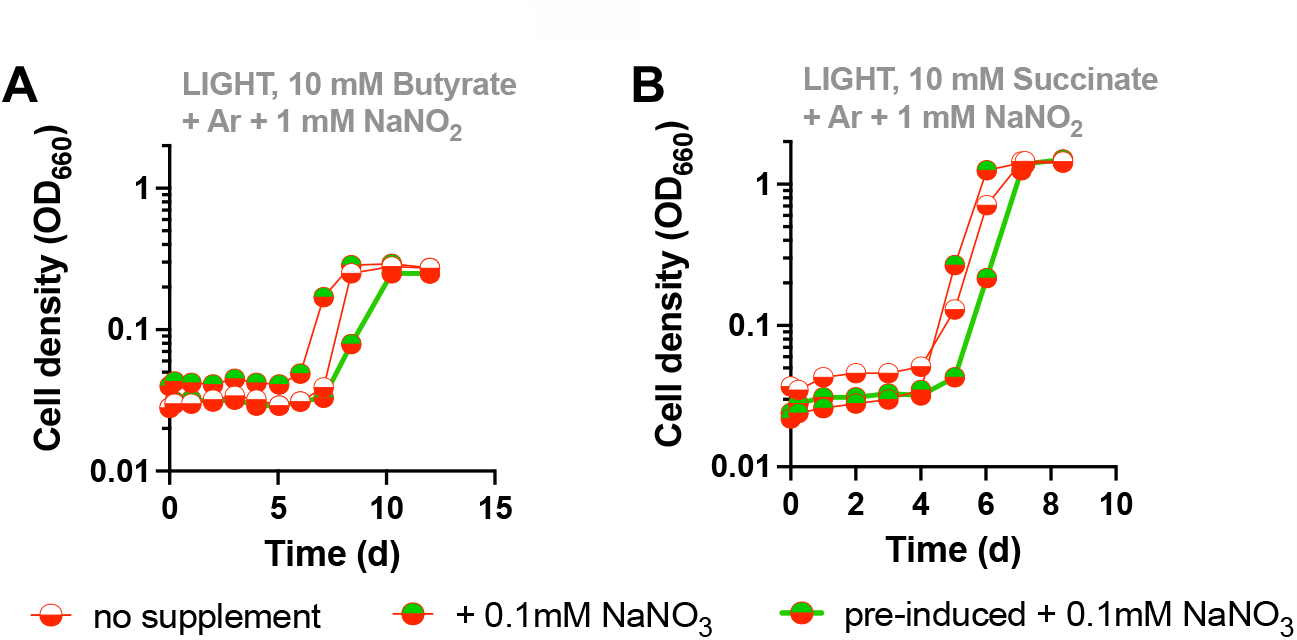
NaNO_3_ does not improve growth trends when NaNO_2_ is present as an essential electron sink. **(A) or as a toxic compound (B)**. Single representatives are shown. Similar trends were observed for three biological replicates.

### NaNO_3_ is required for anaerobic respiration with N_2_O by CGA0092 in the dark

Without access to light, many PNSB can grow chemoheterotrophically via anaerobic respiration. We tested if NaNO_2_ or N_2_O could support chemoheterotrophic growth by CGA0092 in the dark. Acetate and butyrate were chosen as two carbon sources that are metabolized via similar pathways but contain different amounts of electrons (19). Unlike phototrophic conditions (Fig. 2), supplementation with either 0.3 or 1 mM NaNO_2_ did not support observable growth in the dark with either acetate or butyrate within 15 days. In contrast, N_2_O supported growth with either acetate or butyrate but only when NaNO_3_ was also provided (Fig. 7). Because growth was slower with acetate than with butyrate (doubling time ± SD = 88 ± 2 h vs 51 ± 3, respectively), we performed further analyses with butyrate. As during phototrophy (Fig. 3), chemotrophic N_2_ production, culture growth, and butyrate consumption were linearly correlated with the amount of N_2_O provided (Fig. 7C, D). All or nearly all N_2_O was removed while NO_3_^-^ levels remained stable, without NO_2_^-^ production (Fig. 7C).

**Fig. 7.**
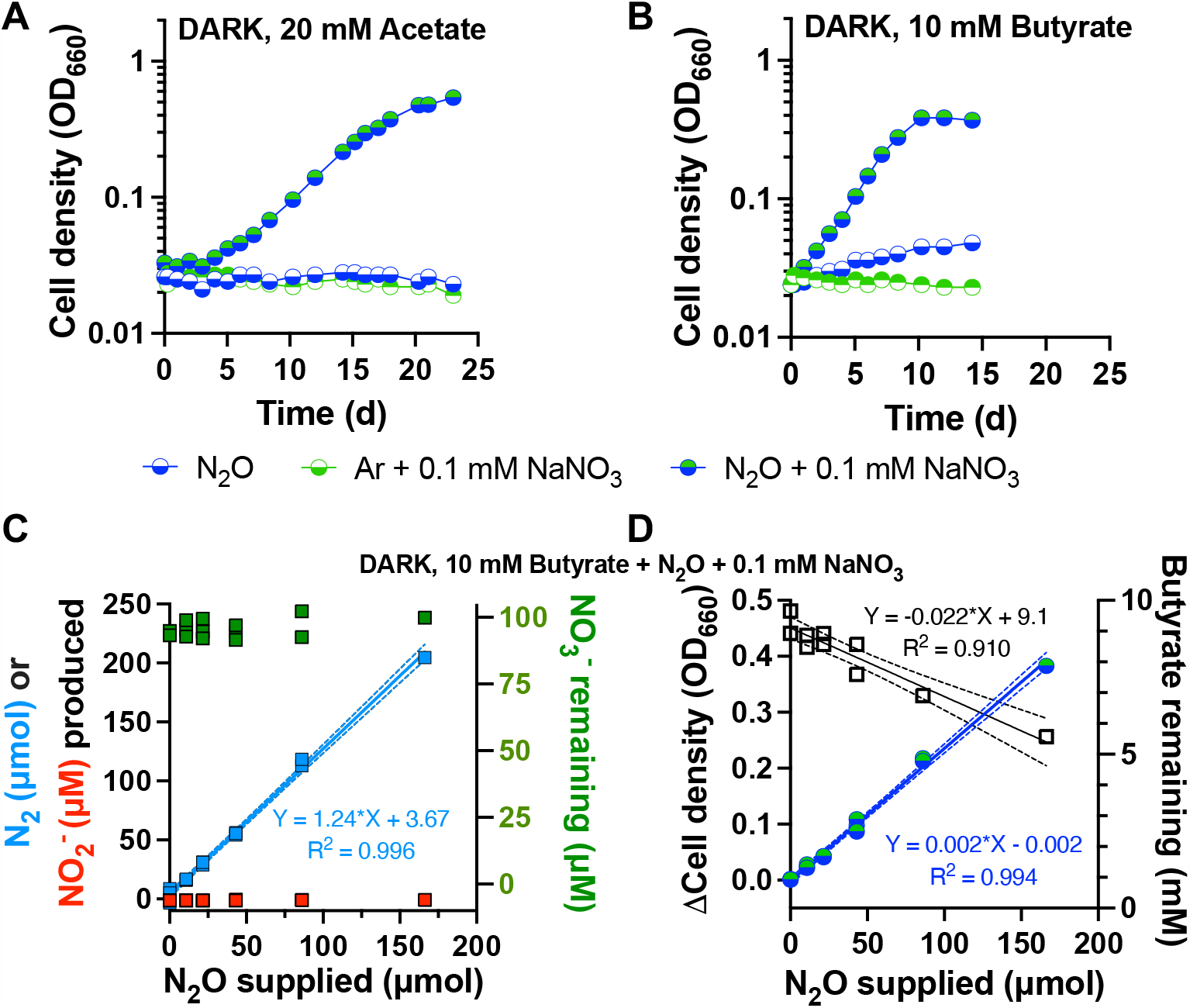
NaNO_3_ is required for N_2_O respiration by CGA0092 in the dark with acetate. **(A) or butyrate (B)**. Single representatives are shown. Similar trends were observed for three biological replicates. Cultures had a 100% Ar headspace or 100% N_2_O headspace as indicated. **C, D**. Changes in N_2_, NO_3_^-^, cell density, and butyrate in N_2_O-limited cultures. Measurements were taken at inoculation and when the maximum cell density was reached. Each point represents a single independent culture. Most or all N_2_O was removed by stationary phase. Linear regression for NO_3_^-^ (**C**) gave a slope that was not significantly different from zero (p = 0.11).

The chemotrophic growth rate and growth yield with butyrate was 24% and 17%, respectively, of those observed under phototropic N_2_O-reducing conditions (Table 1). However, the specific rate of N_2_O reduction was 1.4-fold higher under chemotrophic conditions (Table 1), suggesting that the rate of N_2_O reduction needed to support electron balance under phototrophic conditions is less than that possible when N_2_O reduction is needed for energy transformation. In agreement with the lower growth yield, the N_2_O product yield was 3.3-fold higher under chemotrophic conditions (Table 1), indicating that more electrons from butyrate were directed to energy transformation compared to biosynthesis during chemotrophic growth.

**Table 1.** Growth and metabolic parameters from N_2_O-reducing conditions with butyrate.

| Strain, growth condition | Sp. growth rate (d<sup>-1</sup>) | Doubling time (d) | Sp. N<sub>2</sub>O reduction rate (fmol/CFU/d) | Sp. NO<sub>2</sub><sup>-</sup> reduction rate (fmol/CFU/d) | Growth yield (CFU/pmol N<sub>2</sub>O) | Growth yield (CFU/pmol butyrate) | N<sub>2</sub>O product yield (mol/mol butyrate) |
| --- | --- | --- | --- | --- | --- | --- | --- |
| CGA0092, light | 1.36 ± 0.09 | 0.51 ± 0.04 | 1877 ± 182 | ND | 70 ± 5 | 90 ± 10 | 1.3 ± 0.1 |
| CGA0092, dark | 0.32 ± 0.02 | 2.20 ± 0.16 | 2645 ± 221 | ND | 12 ± 0 | 50 ± 10 | 4.3 ± 0.8 |
| SB1003, light | 1.32 ± 0.18 | 0.53 ± 0.07 | 1503 ± 229 | 5 ± 3 | 87 ± 6 | 105 ± 5 | 1.6 ± 0.3 |
| SB1003, dark | 0.79 ± 0.16 | 0.87 ± 0.02 | 6332 ± 1348 | 5 ± 4 | 13 ± 0 | 75 ± 10 | 5.9 ± 0.8 |

### NaNO_2_ is required for phototrophic N_2_O reduction by *R. capsulatus* SB1003

We wondered if the requirement for non-catalyzable denitrification intermediates for N_2_O utilization was specific to *R. palustris* or was also true in other partial denitrifiers. To examine this possibility, we turned to *R. capsulatus* SB1003, which stood out as an easily cultivatable and phylogenetically distant PNSB that is annotated to only have N_2_O reductase (10) (Fig. 8A).

**Fig. 8.**
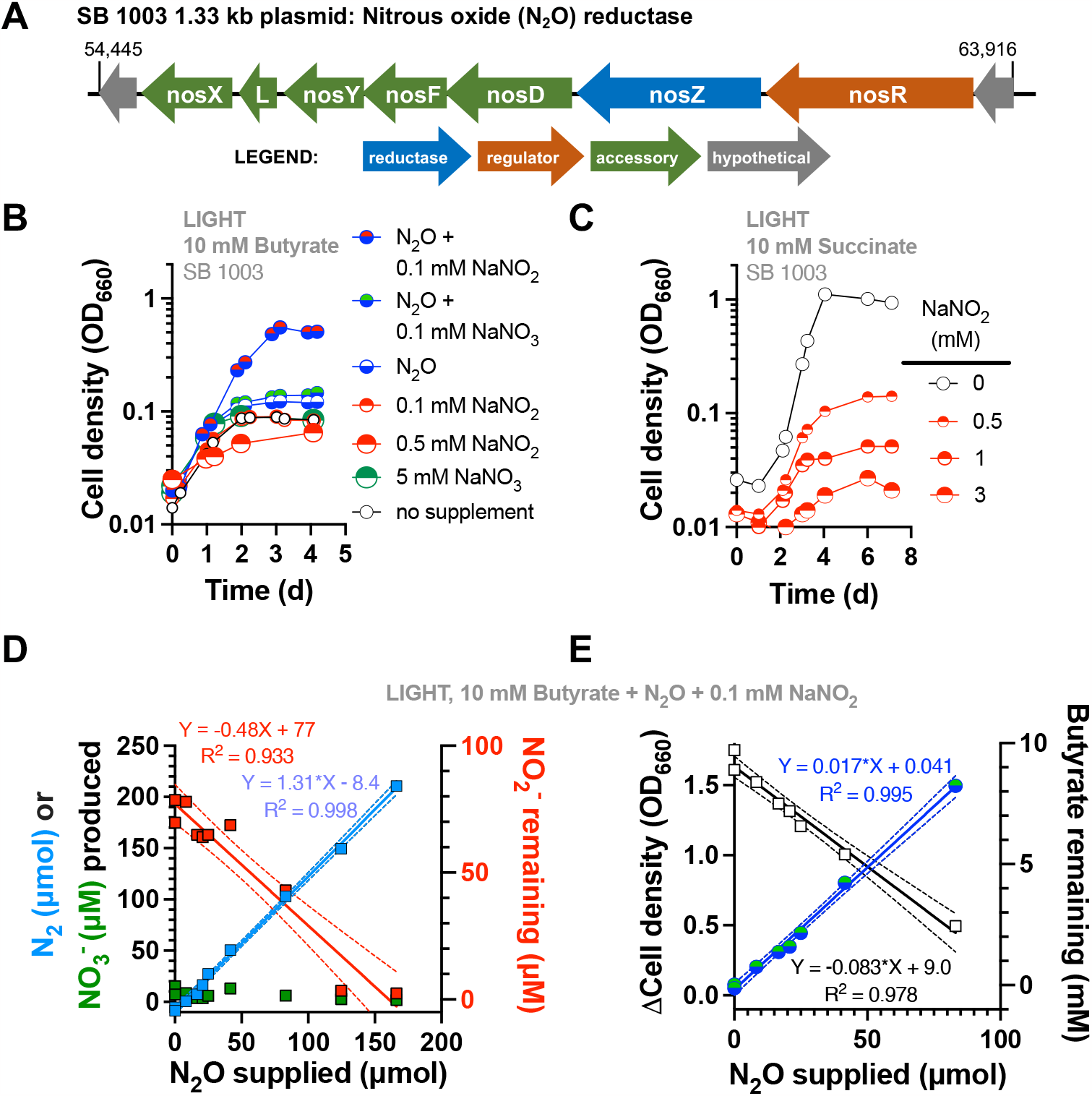
NaNO_2_ is required for photoheterotrophic N_2_O reduction by *R. capsulatus* SB1003. **A**. Plasmid location of predicted N_2_O reductase genes (*nos*). Numbers indicate nucleotide positions. **B**. Phototrophic growth with butyrate and various denitrification intermediates. Single representatives are shown. Similar trends observed for three biological replicates. Cultures had a 100% Ar headspace unless N_2_O is indicated (100% N_2_O). **C**. Phototrophic growth with succinate and various NaNO_2_ concentrations to determine the toxicity limit. Single representatives were surveyed. **D, E**. Changes in N_2_, NO_3_^-^, cell density, and butyrate in N_2_O-limited cultures. Measurements were taken at inoculation and when the maximum cell density was reached. Each point represents a single independent culture. Samples were diluted in cuvettes where necessary to ensure linear correlation between OD and cell density. Most or all N_2_O was removed by stationary phase. Linear regression for NO_2_^-^ (**D**) gave a slope that was significantly different from zero (p-value = 0.0002).

Using PSI-BLAST, we built upon past analyses (10) and confirmed that SB1003 does not have genes with significant similarity to known assimilatory and dissimilatory NO_3_ and NO_2_ reductase genes (Table S1). As predicted, phototrophic growth of SB1003 on butyrate was not supported by NaNO_3_ or NaNO_2_ (Fig. 8B), although our ability to assess the latter was limited by the sensitivity of SB1003 to NaNO_2_ concentrations > 0.5 mM (Fig. 8C). Similar to what was observed for *R. palustris*, N_2_O alone did not support phototrophic growth of SB1003 (Fig. 8B). However, supplementation with 0.1 mM NaNO_2_, but not NaNO_3_, led to phototrophic growth on butyrate with N_2_O (Fig. 8B). Also, N_2_ production, growth, and butyrate production were linearly correlated with the amount of N_2_O provided (Fig. 8D, E), with all or nearly all N_2_O removed. However, unlike with *R. palustris*, levels of the stimulating compound, in this case NO_2_^-^, were not stable. NO_2_^-^ concentration declined with a roughly linear correlation to the amount of N_2_O provided (Fig. 8D). NO_3_^-^ concentrations remained close to zero (Fig. 8D), suggesting that NO_2_^-^ was reduced, rather than oxidized. The specific rate of N_2_O reduction was 300-times higher than that of NO_2_^-^ reduction (Table 1). This disparity suggests that NO_2_^-^ removal was likely due to a promiscuous enzyme activity or a growth-correlated abiotic factor rather than due to an unannotated bonafide NO_2_^-^ reductase.

Values show averages ± 95% CI. Doubling time = ln2/growth rate. Specific (Sp) reduction rates were determined by multiplying the growth rate by the slope of a linear regression of product vs cell density (30, 31) from N_2_O-limited cultures. Colony forming units (CFU) were obtained using a conversion factor of 1 OD_660_ = 5 x 10^8^ CFU/ml (32). Product yields were determined by linear regression using measurements of N_2_O, NO_2_ ^-^, and butyrate from N_2_O-limited cultures.

### NaNO_2_ is required for anaerobic respiration with N_2_O by SB1003 in the dark

SB1003 could also respire N_2_O in the dark with butyrate, but again only when NaNO_2_ was also present (Fig. 9A). Most or all N_2_O was converted to N_2_ when limiting amounts of N_2_O was provided in the presence of 0.1 mM NaNO_2_ (Fig. 9B). N_2_O supplied also showed linear correlation with culture growth and butyrate consumption (Fig. 9C). Although only a small amount, some NO_2_^-^ was likely removed during N_2_O reduction because the linear correlation of NO_2_^-^ levels with N_2_O supplied was negative and significantly different from zero (Fig. 9B). The specific NO_2_^-^ reduction rate was 3-orders of magnitude slower than the specific N_2_O reduction rate (Table 1), again suggesting that the activity was not associated with a canonical denitrification reaction. Similar to *R. palustris*, the SB1003 chemotrophic growth rate and growth yield were lower than those in phototrophic conditions, and more electrons in butyrate were diverted to N_2_O reduction compared to biosynthesis (Table 1). However, SB1003 appears to be capable of a 2.4-fold higher specific N_2_O reduction rate, which is likely behind the proportionately higher chemotrophic growth rate compared to *R. palustris* CGA0092 (Table 1).

**Fig. 9.**
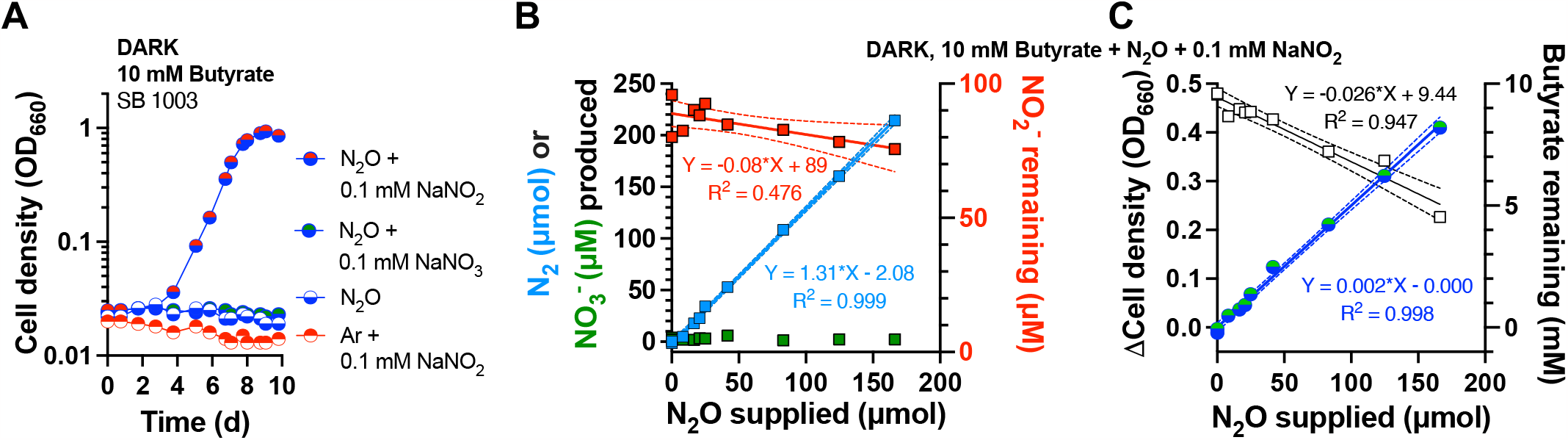
NaNO_2_ is required for N_2_O respiration by SB1003 in the dark. **A**. Chemotrophic growth of SB1003 on butyrate with N_2_O as an electron acceptor. Single representatives are shown. Similar trends were observed for three biological replicates. Cultures had a 100% Ar headspace unless N_2_O is indicated (100% N_2_O). **B, C**. Changes in N_2_, NO_3_^-^, cell density, and butyrate in N_2_O-limited cultures. Measurements were taken at inoculation and when the maximum cell density was reached. Each point represents a single independent culture. Most or all N_2_O was removed by stationary phase. Linear regression for NO_2_^-^ (**B**) gave a slope that was significantly different from zero (p-value = 0.0097).

## Discussion

Here we verified that *R. palustris* CGA0092 and *R. capsulatus* SB1003 are partial denitrifiers, with each being capable of respiring the nitric oxides predicted from their genome annotations. We observed that these bacteria can only reduce N_2_O when supplied with other denitrification intermediates. Most importantly, nitrogen oxides required for N_2_O reduction include those outside of each organism’s partial denitrification repertoire, that is, NO_3_^-^ in the case of CGA0092 and NO_2_^-^ in the case of SB1003.

### NO_3_^-^ as a non-catalyzable inducer of *nos* gene expression

Using a LacZ reporter under control of an *R. palustris nosR* promoter, we found that N_2_O reduction is induced by NO_3_^-^ and NO_2_^-^, at least in part, at the level of gene expression (our construct likely captures transcriptional and translational regulatory features). The level of induction was low compared to the 10-fold increase seen with a *B. diazoefficiens nosZ–lacZ* reporter (24, 33). One possible explanation for the discrepancy is our use of a *nosR* promoter*–lacZ* fusion. NosR is a required regulatory protein for nitrous oxide reductase activity (34, 35) that is typically encoded with little to no intergenic region between *nosR* and *nosZ* (Fig. 1). While the *nosR* promoter drives *nosZ* expression in some bacteria (23), in other bacteria *nosZ* expression can occur from separate, and sometimes multiple, transcriptional start sites (25, 36). Substantial work beyond the current study will be necessary to decipher the transcriptional and post-transcriptional regulatory mechanisms governing *R. palustris nos* genes. However, our findings clearly indicate that NO_3_^-^ or NO_2_^-^ are required for N_2_O reduction by *R. palustris* CGA0092, and that they play a role as inducers of *nos* gene expression.

### Modification of NO_2_^-^ and NO_3_^-^?

A caveat to the observed activation of N_2_O reduction activity by NO_2_^-^ and NO_3_^-^ is that some promiscuous biotic or abiotic transformation could be necessary to generate the inducing/activating molecule. We hypothesize that a promiscuous enzyme activity led to the slow NO_2_^-^ removal in SB1003 cultures (Fig. 8, 9). One possible candidate is sulfite reductase (CysIJ; RCAP_rcc01594 and 03007), which bears homology to assimilatory nitrite reductase. In *E. coli*, CysIJ can convert NO_2_ ^-^ to NH_4_ ^+^ 1.7-times faster than sulfite reduction but with a 200-fold lower affinity for NO_2_^-^ compared to sulfite (k_m_ = 0.8 mM NO_2_^-^, 8-times above the concentration used in our SB1003 experiments) (37). If CysIJ was responsible for NO_2_^-^ removal, then NO_2_^-^ was likely the molecule activating N_2_O reduction because CysIJ would convert NO_2_^-^ to NH_4_ ^+^, which was already present at mM concentrations in the growth medium. However, there could be another enzymatic or spontaneous activity reducing NO_2_^-^ to NO as a separate inducing molecule. For CGA0092, levels of N_2_O reductase-inducing NO_3_^-^ were stable. However, it is still possible that some of the 100 µM NO_3_^-^ was converted below our detection limit to NO_2_^-^ and/or NO. These molecules can induce N_2_O reductase in other organisms like *P. aeruginosa*, although they are typically applied at µM or mM levels (23).

### Possible regulators of N_2_O reduction

Our work calls for future investigation into the regulatory mechanisms controlling N_2_O reduction in CGA0092 and SB1003. However, we can speculate on the regulatory proteins involved based on genome annotations. In considering NO_3_^-^ as an inducing molecule in CGA0092, NasTS stands out as a candidate. In *Bradyrhizobium diazoefficiens*, NasTS controls the transcriptional activation of *nos* genes (24, 25). CGA0092 has genes with significant sequence identity to *B. diazoefficiens* NasTS (77 and 66%, respectively; Fig. 1B). NnrR (Fig. 1B) could also be involved in regulating N_2_O reductase, though more likely in response to NO (1, 23, 38-40).

NO_2_^-^ could also activate N_2_O reduction in SB1003 via an NnrR-like regulator. SB1003 has several CRP/Fnr-family transcriptional regulator genes encoded in its chromosome with >25% amino acid identity to denitrification regulators like *P. denitrificans* FnrP (RCAP_rcc02493; 74% identity) and *Pseudomonas aeruginosa* Dnr/NnrR (e.g., RCAP_rcc00107; 36% identity). One or more of these regulators could be involved in regulating N_2_O reductase, though more likely in response to NO generated biotically or abiotically from NO_2_^-^ (23).

### Denitrification inventories should consider both reductases and regulators

Denitrification gene inventories are notoriously inconsistent with organismal phylogeny; it is common for one species to carry more or less denitrification genes than a close relative (8-10, 12, 41, 42). This inconsistency is also true for strains of *R. palustris* and *R. capsulatus* (20, 43-46). In some cases, horizontal gene transfer (HGT) could explain the phylogenetic discrepancies. The location of the *nos* operon on a plasmid in SB1003 is a straight-forward example of HGT (Fig. 8A). However, HGT cannot explain many of the chromosomal phylogenetic discrepancies. Phylogenetic analyses have suggested the involvement of other factors like gene duplication and divergence, lineage sorting (41), and gene loss (12). Gene loss could be advantageous in communities where there are redundant denitrification functions. As proposed in the Black Queen Hypothesis (47), gene loss can occur when the cost of producing something outweighs the benefit of obtaining it from a neighbor. In this case, the cost of a full denitrification pathway could drive loss of denitrification genes if sufficient energy can be obtained by using a denitrification intermediate released by a neighbor (12). Benefits of partial denitrification pathways have been demonstrated using a synthetic community, though the benefits stemmed more from NO_2_^-^ detoxification than energy savings (48).

Inventories of denitrification regulators have not received the same level of phylogenetic scrutiny as the reductases. Such analyses would be complicated by the fact that phylogenetically similar regulators can regulate different genes (49). However, regulator inventories are likely an important determinant of reductase inventories because improper regulation could influence maintenance or loss of a reductase gene. Regulator inventories also raise questions about the evolutionary histories of denitrification genes. For example, if NasTS is required for expression of the *nos* operon in *R. palustris* CGA0092, were both regulator and reductase genes serendipitously acquired at the same time as separate DNA molecules by HGT or were they acquired together as a single DNA molecule and then physically separated through genome rearrangements (Fig. 1B)? Alternatively, perhaps the common ancestor to CGA0092 and *B. diazoefficiens* USDA110 was capable of complete denitrification and CGA0092 lost nitrate reductase genes but retained the native regulatory network that was responsive to NO_3_^-^. Both genera clade together within the Nitrobacteraceae family but the CGA0092 genome is 3.4 Mb smaller than that of USDA110 and in each case the NasTS and the *nos* genes are separated by large stretches of chromosome (∼ 2Mb in CGA0092 and ∼5Mb in USDA110). Regulator inventory might also support a community role for partial denitrifiers. The regulation of N_2_O reductase by NO_3_^-^ and NO_2_^-^ in this study could suggest that CGA0092 and SB1003 are primed to sense signals by denitrifying partners.

Our findings suggest that within communities of partial denitrifiers, nitric oxides are not only cross-fed metabolites but also important regulatory molecules. The requirement of these molecules for N_2_O reduction is an important consideration in efforts to mitigate greenhouse gas emissions from agricultural soils, which is the largest source of N_2_O emissions (6). Given that agricultural soils are fertilized with NO_3_^-^, we do not anticipate a shortage of NO_3_^-^ in those environments. However, our findings expose a potential pitfall in overlooking the capacity of *nosZ*-harboring bacteria to reduce N_2_O, if unanticipated inducing molecules are omitted from lab cultures.

## Methods

### Strains

*R. palustris* CGA0092 is a chloramphenicol-resistant type strain derived from CGA001 and differs from CGA009 by a single nucleotide polymorphism (13, 14). The Calvin cycle mutant Δ*cbbLSMP*::km^R^ (ΔCalvin, CGA4008) was constructed by deleting *cbbLS*, encoding ribulose-1,5-bisphosphate carboxylase (Rubisco) form I, in a previously described mutant lacking Rubisco form II (Δ*cbbM*; CGA668; (26)) via introduction of the suicide vector pJQΔ*cbbLS* (50) by conjugation with *E. coli* S17 as described (50, 51). The gene encoding phosphoribulokinase, *cbbP*, was then deleted in the resulting strain (Δ*cbbLSM*; CGA4006) by introducing the suicide vector pJQΔ*cbbP*::km^r^ (50), as above, to generate the Δ*cbbLSMP*::km^R^ strain, CGA4008. The elimination of three genes unique to the Calvin cycle greatly decreases the odds of enriching for suppressor mutations. All strain genotypes were verified by PCR and Sanger sequencing. CGA4011 is a NifA* derivative of CGA4008 that has constitutive nitrogenase activity/H_2_ production (50). *R. capsulatus* SB1003 was provided courtesy of Carl Bauer (Indiana University).

CGA4070 was derived from CGA0092 for assaying *nosR* promoter activity using a *lacZ* reporter chromosomally integrated upstream of the *nos* gene cluster, a similar strategy as that used in *B. diazoefficiens* (24, 52). Briefly, a 398-nt region upstream of the *nosR* start codon was synthesized in front of *lacZ* and incorporated into pTwist Kan High Copy plasmid by Twist Bioscience (twistdna.com) to create pTwist_PNos-LacZ. CGA0092 was transformed with pTwist_PNos-LacZ by electroporation and plated on photosynthetic medium (PM) agar with 10 mM succinate and 100 µg/ml kanamycin. Colonies were screened for integration by both PCR and by the appearance of blue color when patched to identical agar that also contained 5-bromo-4-chloro-3-indolyl-β-D-galactoside.

### Growth conditions

Strains were routinely cultivated in 10 ml PM in 27-ml anaerobic test tubes. PM is based on described media compositions (53, 54) and contains (final concentrations): 12.5 mM Na_2_HPO_4_, 12.5 mM KH_2_PO_4_, 7.5 mM (NH_4_)_2_SO_4_, 0.1 mM Na_2_S_2_O_3_, 15 µM p-aminobenzoic acid, and 1 ml/L concentrated base (54). Concentrated base contains: 20 g/L nitriloacetic acid, 28.9 g/L MgSO_4_, 6.67 g/L CaCl_2_·2H_2_O, 0.019 g/L (NH_4_)_6_Mo_7_O_24_·4H_2_O, 0.198 g/L FeSO_4_·7H_2_O, and 100 ml/L Metals 44 (55). Metals 44 contains: 2.5 g/L ethylenediaminetetraacetic acid, 10.95 g/L ZnSO_4_·7H_2_O, 5 g/L FeSO_4_·7H_2_O, 1.54 g/L MnSO_4_·H_2_O, 0.392 g/L CuSO_4_·5H_2_O, 0.25 g/L Co(NO_3_)_2_·6H_2_O, 0.177 g/L Na_2_B_4_O_7_·10H_2_O. PM was made anaerobic by bubbling tubes with 100% Ar then sealing with rubber stoppers and aluminum crimps prior to autoclaving. After autoclaving, tubes were supplemented with either 20 mM sodium acetate, 10 mM sodium butyrate, or 10 mM disodium succinate from 100X anaerobic stock solutions. Where indicated, cultures were additionally supplemented with 20 mM NaHCO_3_. SB1003 cultures were also supplemented with 0.1 µg/ml nicotinic acid, 0.2 µg/ml riboflavin, and 1.3 µg/ml thiamine-HCl. NaNO_2_ or NaNO_3_ were added from anaerobic stock solutions to the final concentrations indicated in the text. For conditions with N_2_O, tubes were flushed with 100% N_2_O through a 0.45 µm syringe filter and needle after all liquid supplements were added. A second needle was used for off-gassing. For N_2_O-limited cultures, the indicated volume of filtered gas was added via syringe. Cultures were inoculated with a 1% inoculum from starter cultures grown phototrophically in anaerobic PM with succinate, except for the experiment testing Calvin cycle mutants (Fig. 3C) in which all starter cultures were grown aerobically in 3 ml PM with succinate in the dark. These aerobic conditions were used to accommodate the Δ*cbbLSMP*::km^R^ mutant (CGA4008) that requires an electron sink to grow.

### Analytical procedures

Culture growth was monitored via optical density at 660 nm (OD_660_) using a Genesys 20 spectrophotometer (Thermo-Fisher, Waltham, MA, USA) directly in culture tubes without sampling. For N_2_O-limitted cultures, samples were diluted in cuvettes where specified. Specific growth rates were calculated using OD_660_ values between 0.1 and 1.0 where cell density and OD are linearly correlated. N_2_, N_2_O, and H_2_ were sampled from culture headspace using a gas-tight syringe and analyzed using a Shimadzu GC-2014 gas chromatograph (GC) equipped with a thermal conductivity detector. GC conditions for H_2_ were described previously (56). GC conditions for N_2_ and N_2_O used He as a carrier gas at 20 ml/min, a 80/100 Porapak N column (6’ x 1/8” x 2.1 mm; Supelco) at 170 °C, an inlet temperature of 120°C, and a detector temperature of 155°C with a current of 150 mA. Gas standards were prepared by injecting specific volumes of 1 ATM of pure gasses (41.6 mM based on the ideal gas law and a temperature of 293 K) into a stopper-sealed serum vial of known volume, containing with a few glass beads to aid in mixing. Gas standards were mixed by shaking, sampled with a gas-tight syringe, and then injected at 1 ATM by releasing pressure prior to injection. Pressure was not released prior to injection for culture headspace samples. Syringes were flushed with He prior to each standard and culture injection to minimize contamination with atmospheric N_2_. NO_3_^-^ and NO_2_^-^ were measured using a colorimetric Griess assay kit according to the manufacturer’s instructions (Cayman Chemical). Conversion of NO_3_^-^ to NO_2_^-^ was accomplished via NO_3_^-^ reductase provided with the kit. N_2_, N_2_O, NO_3_^-^ and NO_2_^-^ were measured at the time of inoculation and at stationary phase.

### β-galactosidase reporter assays

*R. palustris* strains were grown to mid-late exponential phase (0.4 - 1.1 OD_660_) with 23 mM sodium acetate and either 0.1 mM NaCl, NaNO_3_, or NaNO_2_, with or without 4 ml N_2_O. Cultures were then chilled on ice and all subsequent processing was carried out between 0 - 4 °C. Cells were harvested by centrifugation, supernatants were discarded, and cells were resuspended in 0.5 ml Z-buffer. Cells were lysed by five 20-s rounds of bead beating at maximum speed using a FastPrep^®^-24 benchtop homogenizer (MP Biomedical), with 5 min on ice between rounds. Cell debris was pelleted by centrifugation and supernatant protein was quantified using Bio-Rad’s Bradford assay kit. Cell lysate (50 µl supernatant) was mixed with 100 µl Z-buffer in wells of a 96-well plate. Reactions were started with the addition of 30 µl of 4 mg/ml ortho-nitrophenyl-ß-galactoside. Formation of *o*-nitrophenol was monitored at 420 nm over time at 30°C using a BioTek Synergy plate reader. Specific activity was determined by linear regression of the initial velocity and normalized for protein concentration.

### Bioinformatics

PSI-BLAST used default parameters except for 500 targets, an expect threshold of 10, a word size of 3, and a PSI-BLAST threshold of 0.005 using the refseq_protein database for bacteria. At least five iterations were run or until no further sequences were found. Accession numbers for the query sequences are in Table S1.

### Statistical analyses

Graphpad Prism v10 was used for all statistical analyses.

## Acknowledgements

This work was supported in part by a National Science Foundation CAREER award (MCB-1749489NSF), the Division of Chemical Sciences, Geosciences, and Biosciences, Office of Basic Energy Sciences, U.S. Department of Energy (DOE), through Grant DE-FG02-05ER15707, the Office of Science (BER), U.S. Department of Energy, through Grant DE-FG02-07ER64482, and the Indiana University College of Arts and Sciences. We are grateful to Doug Rusch, Julia van Kessel, Cristina Landeta, Nick Haas, José Heerdink-Santos, Jillian Lewis, William Rockliff, and Anika Hays for advice, reagents, and media preparation. We are also grateful to the anonymous reviewers for constructive feedback.

